# Loss of SUR1 and Kir6.2 ATP-sensitive potassium channel subunits produces comparable neuronal excitability and morphine antinociception

**DOI:** 10.64898/2026.10.27.761978

**Authors:** Soomin C. Song, Brie Beck, Kayla Johnson, Sunna Mujteba, William A. Coetzee, Amanda H. Klein

## Abstract

ATP-sensitive potassium (K_ATP_) channels are integral membrane proteins that play a crucial role in linking cellular metabolism to neuronal excitability. KATP channels in peripheral nociception and pain emphasize metabolic gating as a determinant of dorsal root ganglia and spinal cord excitability, particularly in pathophysiological states. Activation of K_ATP_ channels reduces excitability and and also contributes to opioid induced analgesia,. K_ATP_ channels are composed of multiple subunits, that are assembled in several subtypes, including Kir6.2/SUR1 which is found predominantly in neurons in the nervous system. It remains unclear whether both Kir6.2 and SUR1 subunits are required to regulate neuronal excitability under naïve conditions and to mediate morphine-induced antinociception. These experiments aim to determine whether neuron-specific loss of either Kir6.2 or SUR1 increases dorsal root ganglion (DRG) neuronal excitability, and attenuates morphine antinociception in mice. We found that loss of SUR1 or Kir6.2 activity or expression both increased DRG excitability in whole cell patch clamp studies. Using DRG cultured from SUR1 and Kir6.2 cKO animals, we also found that loss of either subunit decreased potassium flux induced by diazoxide. Morphine induced antinociception was also attenuated after Cre-mediated recombination in SUR1 and Kir6.2 cKO animals. Chronic morphine administration also decreased Kir6.2 (gene: Kcnj11) RNA signal in mouse DRG, but not spinal cords. Together, these findings suggest that, despite evidence from single-cell sequencing and other datasets indicating potentially distinct expression patterns of SUR1 and Kir6.2, these subunits most likely function cooperatively to form functional K_ATP_ channels under normal physiological conditions.

## Introduction

ATP sensitive potassium (K_ATP_) channels act as metabolic gatekeepers of neuronal excitability in the peripheral and central nervous system. K_ATP_ channels open during periods of low ATP:ADP ratio, where they are involved in hyperpolarizing neurons, and help prevent metabolic overload during stress. Conversely, when cellular ATP is abundant and ADP levels are low, they remain closed, supporting normal excitability. K_ATP_ channels are an octameric protein complex composed of four pore-forming Kir6.x subunits and four regulatory sulfonylurea receptor (SUR) subunits (Mikhailov et al., 2005). Suface trafficking of the receptor requires both subunits, but the SUR subunit, is thought to be crucial for ADP, diazoxide, and PIP_2_ sensitivity (Gribble et al., 1997; Pratt et al., 2011; Zerangue et al., 1999). In the nervous system, the Kir6.2/SUR1 complex is the predominant subtype, particularly in neurons in the peripheral nervous system (Kawano, Zoga, McCallum, et al., 2009). Loss of K_ATP_ channel activity contributes to hypersensitivity in chronic pain models (Chi et al., 2007), while application of K_ATP_ channel agonists, predominately those known to target SUR1-subunits, have been shown to be antinociceptive (Ocana et al., 1996; Welch & Dunlow, 1993) and have analgesic effects in preclinical pain models (Beich et al., 2025). K_ATP_ channels are also involved in downstream signaling of mu-opioid receptors, and agonists enhance the analgesic efficacy of opioids (Afify et al., 2013; Cunha et al., 2010; Rodrigues & Duarte, 2000), and improve opioid tolerance and withdrawal in rodent models (Fisher, Johnson, et al., 2019).

*Kcnj11* and *Abcc8*, which code for the K_ATP_ channel subunits Kir6.2 and SUR1, respectively, are co-localized in mechanosensory neurons in the dorsal root ganglia, including low threshold mechanosensory types in humans (Nguyen et al., 2021). SUR1 mRNA transcripts are also located in nociceptive peptidergic classes of DRG neurons (Kupari et al., 2021) and low-threshold A-delta fibers (Jung et al., 2023). In human spinal cord, *KCNJ11* and *ABCC8* mRNA are expressed in several neuron classes, however, *ABCC8* is expressed in other classes of excitatory neurons, microglia, astrocytes, and oligodendrocytes without *KCNJ11* (Yadav et al., 2023). Similar patterns arise in the human brain, where single cell RNA sequencing indicates that *Kcnj11* and *Abcc8* are found on glutamatergic and GABAergic neurons (Constantino et al., 2025). More specifically, *Abcc8* and *Kcnj11* are expressed in not completely overlapping neuron clusters in the spinal cord that follow the spinothalamic, spinoreticular, and spinomesencephalic tracts (i.e. anterolateral system) (Bell et al., 2023). In the non-overlapping zones, they may function strictly as homomers or with alternative partners (e.g. SUR2A, SUR2B, Kir6.1), establishing unique regional signaling profiles.

K_ATP_ channel structure and function studies indicate that SUR1 is a regulatory subunit, responsible for increasing the open probability of the channel after exposure to agonists and sensitivity to therapeutic drugs, including sulfonylureas (e.g. glibenclamide) that close the channel (Li et al., 2021). Global loss of SUR1 does not greatly increase baseline hypersensitivity to mechanical or heat thresholds (Fisher, Johnson, et al., 2019), but does decrease morphine antinociception in mice (Sakamaki et al., 2021). Loss of Kir6.2 in aged mice results in hyposensitivity to thermal stimuli compared to wild-type controls, and results in some myelinated, but mostly unmyelinated, nerve fiber degeneration(Nakai-Shimoda et al., 2022). These studies provide strong evidence for K_ATP_ channel subunit roles in peripheral nervous system activity, but the global loss of SUR1 or Kir6.2 could alter insulin secretion (Seghers et al., 2000) or glucose tolerance (Seino et al., 2000), which could confound behavioral and electrophysiological studies. It is unknown if Kir6.2 and SUR1 subunits are both required for modulating neuronal excitability under naïve conditions, or if both subunits are necessary for facilitating morphine analgesia. The purpose of these experiments is to determine if loss of Kir6.2 or SUR1 in neurons increases DRG excitability or decreases morphine antinociception. Since K_ATP_ channel stimulation or antagonism affects behavioral outputs to opioid tolerance and withdrawal, we also utilized *in situ* hybridization to determine expression of each subunit independently after chronic opioid administration.

## Methods

### Animals

*Kir6.2*^flox/flox^ (Yang et al., 2016) and SUR1 ^flox/flox^ (Nakamura & Bryan, 2014) and wild-type littermates were bred on a congenic C57BL/6 background. Both male and female 8-12 week old mice were used in all experiments. All mice were group-housed in an Association for Assessment and Accreditation of Laboratory Animal Care International accredited facility on a 12 h light/dark cycle (zeitgeber time or ZT 0, lights on at 7:00 A.M. CST/EST) with *ad libitum* access to food and water. All experiments were approved by the Institutional Animal Care and Use Committee of the University of Minnesota, University at Buffalo, and New York University School of Medicine. Genotyping was performed by polymerase chain reaction (PCR) using the HOTSHOT method of DNA preparation and Quick-Load® Taq 2X Master Mix (New England Biolabs). Genotyping was confirmed using: Kir6.2 (*Kcnj11*) Flox forward primer (5′-CCA GTG CCC CTA TTA CCC TAG C-3′), Kir6.2 Flox reverse primer (5′-GAG TTT GGG CCT GAG CAG CTG ATT C-3′) with an identifying band at 551bp for a WT mouse and at 728bp for a floxed mouse. The SUR1 (*Abcc8*) Flox PCR used forward primer (5′-TGA GAT CGC TGA GGG TAT CC-3′) and reverse primer (5′-GGG CTG TGC ACT GTG AAT AC-3′) with an identifying band at 551bp for a WT mouse and at 728bp for a floxed mouse.

### Viral Delivery

To establish a conditional knockout (cKO) of SUR1 and Kir6.2 in lumbar spinal cord and DRG, intrathecal injections of either AAV9-hSyn-RFP-Cre or AAV9-hSyn-RFP-ΔCre (5 μL containing ∼10^13^ vector genomes, University of Minnesota Viral Vector Core, Minneapolis, MN, United States) were performed using lumbar puncture method (Pflepsen et al., 2019; Vulchanova et al., 2010). Four weeks post viral delivery mice were weighed to determine dosing for behavioral experiments or used for electrophysiology.

### Whole-cell Patch Clamping of DRG Neurons

Adult C57Bl/6J male and female mice were deep anesthetized with 5% isoflurane before cervical dislocation and decapitation. The spinal column was then bisected to expose the spinal cord and DRGs. Individual DRGs were excised from the spine and placed into HBSS. After collection of the DRGs, individual DRGs were fileted and placed in 2:1 of culture media to digestion media. Culture media consisted of DMEM with 10% fetal bovine serum, 1% penicillin-streptomycin, and 30 ng/mL nerve growth factor. Digestion media consisted of 3.125 mg/mL neutral protease and 5 mg/mL collagenase Type 1 in DMEM. DRGs in digestion media were then dissociated in a heated rocker (37°C) for 1 hour. Cells were pelleted and rinsed with culture media before being plated on poly-D-lysine + laminin coated cover slips and left to rest in a cell culture incubator for 1 h – 24 h prior to recording.

Whole-cell patch clamp recordings were performed using the traditional patch clamping technique to measure cell excitability of DRG neurons. Cover slips were transferred to the stage of an upright Olympus BX51WI microscope and superperfused with oxygenated artificial cerebral spinal fluid (ACSF) consisting of (in mM): 124 NaCl, 2.5 KCl, 1.25 NaH_2_PO_4_, 26 NaHCO_3_, 10 D-glucose, 1.5 MgSO_4_ and 2.5 CaCl_2_.

Pipettes of borosilicate glass were manufactured with a DMZ Universal Puller (Zeitz, Germany). Pipette resistances were 4-5 MΩ when filled with an internal solution containing (in mM): 127 K-gluconate, 8 KCl, 10 phosphocreatine, 10 HEPES, 4 MgATP and 0.3 NaGTP; pH of 7.3 adjusted with KOH; osmolarity 284.5 mOsm. Recordings were made using an Axopatch 200B amplifier (Molecular Devices), digitized (Axon Digidata 1550A; Molecular Devices) and recorded with Clampex 10.7 software (Molecular Devices). The currents were filtered at 2 kHz (low pass, 8-pole Bessel response) and digitized at 10 kHz. Whole cell capacitance and series resistance were not compensated. Recordings were not adjusted for the liquid junction potential, which was calculated to be ∼9.4 mV.

The frequency/current relationship (F/I curve) was measured by applying a series of 1 second current steps to hyperpolarize or depolarize the cell to initiate action potentials. The starting, ending and current step amplitude was adjusted for each cell to elicit action potentials on a cell to cell basis.

### Mechanical and Thermal Paw Withdrawal Testing

Animals were acclimated to one of the testing platforms for three consecutive days before experimentation. Mechanical nociception was measured using fine blunt probe pressed on the plantar surface of the hind paw connected to an Electronic Von Frey Anesthesiometer (Model 2390, IITC Life Science, Inc., Woodland Hills, CA) which included an elevated mesh floor with clear acrylic holding areas split into four separate chambers. Radiant heat thermal paw withdrawal was measured on a heated glass platform with acrylic holding areas. A small light source was positioned under the glass, focusing the beam on the plantar surface of the hind paw (Model 290G, IITC). Stimulation was removed when a nocifensive paw withdrawal response was recorded. Baseline measurements were recorded at least five times separately on the left and right hind paws, and the results averaged for each animal. To record morphine-induced antinociception, morphine (Hikma, Catalog # 462-673-01) was administered subcutaneously at ∼45-minute intervals and paw withdrawal thresholds were recorded for each hind paw 30 minutes after each morphine dose (1, 2.5, 5, 10 mg/kg, sc, 100 uL, in saline).

### DRG Extraction and Culture for Biochemical Analysis and Microscopy

Animals were euthanized by an overdose of pentobarbital sodium solution or carbon dioxide as these methods are approved by the American Veterinary Medical Association. Lumbar dorsal root ganglion (DRG) and spinal cord removal from the spinal column were performed using a laminectomy. Spinal cords were flash frozen and stored at -80°C for RNA extraction and qPCR. DRGs were washed in Hank’s Buffered Salt Solution (Corning, 55-022-PB) followed by mincing with small spring scissors. DRG tissues were subjected to enzymatic digestion by a brief incubation with a 37° C papain solution which was composed of 1mg L-cysteine, 3μL saturated NaHCO_3_, 43μL of 20 U/mg papain (Worthington Biochemical Corporation, LS003126) in 1mL of HBSS. After centrifugation and supernatant removal, a second enzymatic digestion was performed at 37° C using a collagenase solution composed of 2mg collagenase (Worthington Biochemical Corporation, LS004176) in 2 mL HBSS. After centrifugation and collagenase removal, DRG were resuspended in 1 mL of complete media containing Minimum Essential Medium Eagle with Glutamine (Fisher Scientific, MT10010CV), 10% horse serum (Fisher Scientific,16-050-122), 1% penicillin and streptomycin (Fisher Scientific, MT30002CI) and 1% vitamin solution (Fisher Scientific,11-120-052). The cell suspension was triturated using fire polished glass Pasteur pipettes. After centrifugation and supernatant removal, DRG were resuspended in enough complete media for plating unto poly-D-lysine (Fisher Scientific Company, 50-165-6956) coated 35 mm glass bottomed plates (Cellvis, D35141.5N). Dishes were placed in a 37° C incubator (humidified with 5% CO_2_) until imaging the following day.

### DRG Live Cell Imaging

Complete media was removed and replaced with 1 mL fresh media and 1 mL of loading dye from the Brilliant Thallium Flex Assay (Ion Biosciences, 11000-100). The plates were then returned to the 37° C incubator for one hour before imaging. A thallium stimulus solution was prepared with or without 5 µL 10 mM diazoxide (in DMSO) to a final concentration of 100 µM in 500 uL (VWR, D5402-1G) to be used during imaging. The plates were imaged using a Leica SP8 STED confocal microscope at 20X to visualize GFP and RFP signal. Time-lapse imaging for GFP occurred over a ten-minute period including 30 seconds of baseline fluorescence before addition of the thallium stimulus solution via perfusion. The resulting EGFP time lapse and corresponding RFP images were analyzed using ImageJ to determine the percent signal intensity pre and post thallium solution addition.

### RNA Extraction and cDNA synthesis

RNA extraction of collected spinal cords was performed by tissue homogenization with glass mortar and pestle followed by the Direct-zol™ RNA Miniprep Kit (Zymo Research, R2052) according to manufacturer instructions. Incubation with DNase I (Zymo Research, catalog # E1011-A) was added to reduce genomic DNA carryover. RNA purity was measured using a nanodrop (NanoDrop 2000c, Thermo Scientific,) or Qubit fluorometer (033216, Invitrogen by Life Technologies). RNA was reverse transcribed into cDNA using the High-Capacity cDNA Reverse Transcription Kit (ThermoFisher Scientific, 4368813). Each sample was loaded with 500 ng of RNA into a reaction containing either the reverse transcriptase enzyme or molecular grade water. All products were placed in a thermal cycler: 1) 25°C for ten minutes, 2) 37°C for 120 minutes, 3) 85°C for five minutes, 4) Hold at 4 °C. After the cycles were complete, the cDNA reaction (20uL) was combined with 180µL molecular grade water and stored at -80°C.

### qPCR

Quantitative Polymerase Chain Reaction (qPCR) was performed using SYBR® Green Supermix (Bio-Rad, 1725270) according to manufacturer instructions. The qPCR reactions included of 2 uL of cDNA (reverse transcriptase positive or reverse transcriptase negative) and 8 uL o f a master mix 0.5 µL forward and reverse primer, 2 µL mbH2O [Fisher Scientific Company LLC, catalog # MT46000CV], and 5 µL SYBR® Green Supermix [Bio-Rad, catalog # 1725270], multiplied by the number of samples. 8 µL of the master mix was pipetted into either a 96 or 384 well plate, followed by 2 µL of reverse transcriptase positive or reverse transcriptase negative cDNA. Amplification was performed on a Bio-Rad CFX96 or a CFX Opus 384 qPCR System (BioRad) cycled at: 95°C for 5 min, followed by 45 cycles of: 94°C for 30 s, 60°C for 10 s, 72°C for 20 s. A melting curve from 65°C to 95°C at increments of 0.5°C for 2 seconds was performed as a post-amplification quality control step. All samples and standards were run in triplicate. A six-point standard curve was generated from a serial dilution of DNA products using specific primers (Table 1). The quantification was performed using BioRad Maestro software (Version 2.3).

**Table 1.**
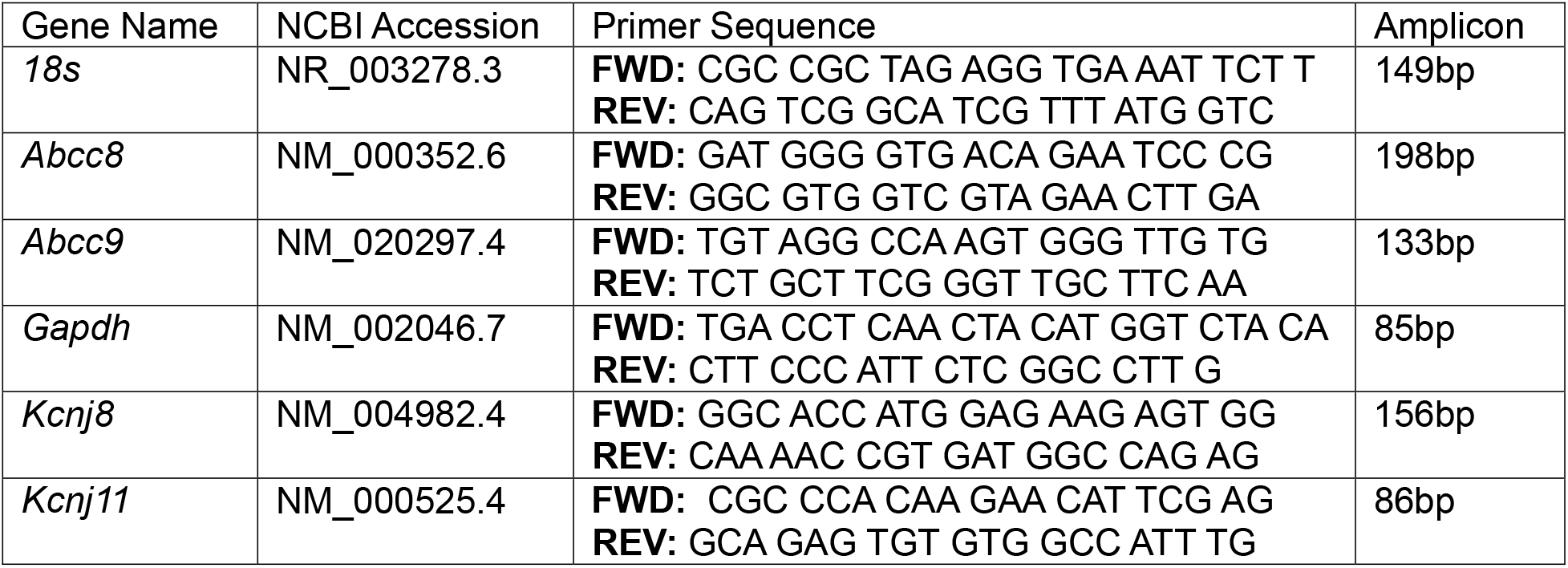
List of primers used in this study.

### In situ hybridization

Mouse lumbar DRG and spinal cords were prepared for in situ hybridization using the RNAscope multiplex version 2 assay (Advanced Cell Diagnostics, Newark, CA) according to manufacturer instructions and technical support. Tissue sections were cut for DRGs and spinal cords at 10 uM and 25 uM, respectively on a cryostat and stored at 80 °C. Slides were transferred to cold (4 °C) 10% normal buffered formalin for 15 minutes. After rinsing in phosphate buffered saline (PBS), the tissues were then dehydrated in a series of ethanol dilutions for five minutes each. Boundaries were drawn around each section using a hydrophobic barrier pen after briefly drying each slide and allowed to set for five minutes. Slides were placed in a humidity-controlled tray and 1 drop of Hydrogen Peroxide was added to each section and allowed to incubate for 10 minutes. Sections were rinsed twice with distilled water for 15 seconds and tapped dry. Protease IV reagent was added to each section and allowed to incubate for 30 minutes at room temperature. After rinsing with PBS, each slide was then placed in a humidity control tray at 40 °C for two hours with a probe mixture for Channel 1 (*Abcc8*) and Channel 2 (*Kcnj11*, 50:1 dilution) pipetted onto each section. After incubation in wash buffer for two minutes, two drops of RNAScope Multiplex Amp1 reagent were added to each section and incubated for 30 minutes at 40 °C. This process was repeated with RNAScope Multiplex Amp2 and Amp3, with a step for rinsing in wash buffer between each incubation. One to two drops of the RNAscope Multiplex FL v2 HRP-C1 reagent was added to each section, incubated for 15 minutes at 40 °C and rinsed with wash buffer. Immediately after, ∼30 uL of OpalTM 520 (Akoya Biosciences) was applied to each section incubated for 30 minutes at 40 °C. After rinsing with wash buffer, RNAscope Multiplex FL v2 HRP Blocker was added to each tissue section and allowed to incubate for 15 minutes at 40 °C. This process was repeated with RNAscope Multiplex FL v2 HRP-C2 reagent and OpalTM620. Following the HRP blocker incubation and rinsing with was buffer, slides were air dried, and cover-slipped with Flouromount G mounting medium with DAPI (Southern Biotech).

### Data Analysis

Patch-clamp recordings were analyzed using custom written code in MATLAB (Mathworks). The code is available upon request. Peak frequencies were compared using either a paired T-test or one-way ANOVA with Prism 9 (Graphpad), when appropriate. The resting membrane potential (RMP) were compared with a paired T-test. A phase-plane analysis on spontaneous action potentials was performed to compare the spike characteristics between groups using one-way ANOVA. F/I curves were determined from the number of spikes generated during a 1 s depolarizing steps. The linear portion of the F/I curves, determined individually for each cell, were compared by fitting a linear regression and compared using an ANCOVA. All error bars are standard error of the means.

## Results

### Activity of mouse DRG modulated by pharmacological modulation of K_ATP_ channels

We conducted whole-cell patch clamping with cultured mouse DRG, which exhibited robust electrical behavior (Figure 1). Glibenclamide is potent blocker of SUR1-subtype K_ATP_ channels with an EC_50_ of ∼2–30nM, while the affinity for SUR2 subtypes is at least 10-25-fold lower(Meyer et al., 1999). Under current clamp conditions, typical action potential firing occurred when DRG neurons were depolarized with current injection (Figure 1A). Diazoxide, which opens SUR1-containing K_ATP_ channels even at low ADP intracellular concentrations(D’Hahan et al., 1999; Dabrowski et al., 2003), inhibited excitability of the DRG neurons under these conditions(Figure 1B). Cells were also less excitable after application of CL-705G. a novel Kir6.2-selective agonist(Gando et al., 2023) (Figure 1C). To analyze this effect in more detail, frequency/current (F/I) curves were constructed. Spike frequency during depolarization of DRG neurons was increased by glibenclamide (Figure 1D) and decreased by diazoxide and CL-705G (Figure 1E,F). For example, a depolarizing current of 300 pA led to a spike frequency of ∼2.5 Hz without drug, whereas the spike frequency was ∼1.5 Hz in the presence of diazoxide (Figure 1E). Peak frequencies were not significantly altered by of any of the compounds (Figure 1G-I). Glibenclamide did not change the resting potential (Figure 1J), but diazoxide and CL-705G both led to resting membrane hyperpolarization (Figure 1K,L). These data indicate that activation of SUR1 or Kir6.2 subunits decrease excitability of peripheral neurons.

**Figure 1.**
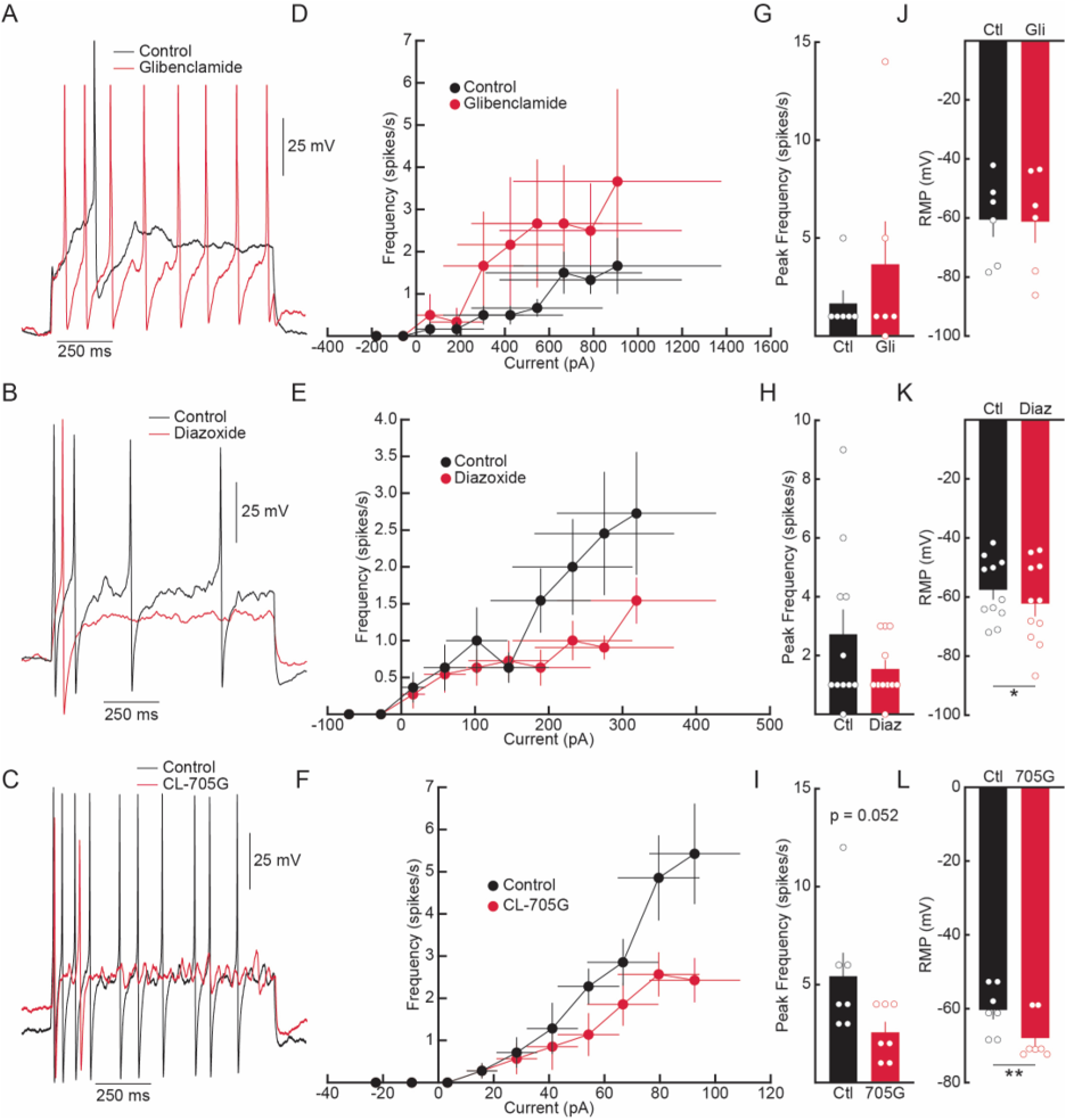
Modulation of K_ATP_ channels in DRG neurons effect cell excitability. A-C) Examples of evoked action potentials with a depolarizing current application under control and modulators of K_ATP_ (1 μM glibenclamide), SUR1 subunit activator (100 μM diazoxide) and K_IR_6.2 activator (100 μM CL-705G). D-F) Mean F/I curves of DRG neurons under control and drug application. D) The slopes of the linear portion were determined to be: control, y = 0.0022 x – 0.29; glibenclamide, y = 0.0026 x + 0.98. The y-intercepts were found to be significantly different (n = 6, F = 6.216, p = 0.0151) implying a change in rheobase after glibenclamide application. E) The equation of the linear portion of the F/I curves were determined to be: control, y = 0.0087 x – 0.072; diazoxide, y = 0.0031 x + 0.26. The slopes were found to be significantly different (n = 11, F = 4.260, p = 0.0407). F) Simple linear fits were y = 0.089x – 2.53 for control and y = 0.032x – 0.43 after CL-705G application. The y-intercepts were significantly different (n = 7, F = 4.150, p = 0.0466). G-I) Comparison of peak frequencies achieved during the current application protocols. I) CL-705G application trended to reduce the peak frequency but was not significant. J-L) RMP of DRG neurons under control and drug application. K) Diazoxide application significantly hyperpolarized the RMP by paired T-test. L) Application of CL-705G also hyperpolarized the RMP significantly, by paired T-test. Error bars are SEM. Open circles are individual data points. *p < 0.05, **p < 0.01.

### Loss of SUR1 or Kir6.2 expression increases excitability of mouse DRG

The roles of Kir6.2 or SUR1 to excitability were further investigated with conditional knockout experiments by intrathecal viral delivery of AAV9-hSyn-RFP-Cre (or AAV9-hSyn-RFP-ΔCre as a genetic control) in *Kir6.2*^flox/flox^ or SUR1 ^flox/flox^ mice (see Methods for details). Local knockout of SUR1 resulted in enhanced excitability of DRG neurons upon depolarization (Figure 2A) compared to wildtype (WT) or ΔCre negative controls. Note that ΔCre injection did not change frequency/current curves and peak frequencies when compared to WT (Figure 2B). Conditional knockout of Kir6.2, also increased the frequency of action potentials firing during depolarization, and the slopes of the F/I curve were significantly different (p = 0.0032; Figure 3). Overall, the pharmacological and the genetic approaches to modulate K_ATP_ channel subunit activity and expression, agree with previous reports that suggest an inverse relationship between activity/expression of K_ATP_ channels and neuronal excitability (Kawano, Zoga, McCallum, et al., 2009; Zoga et al., 2010).

**Figure 2.**
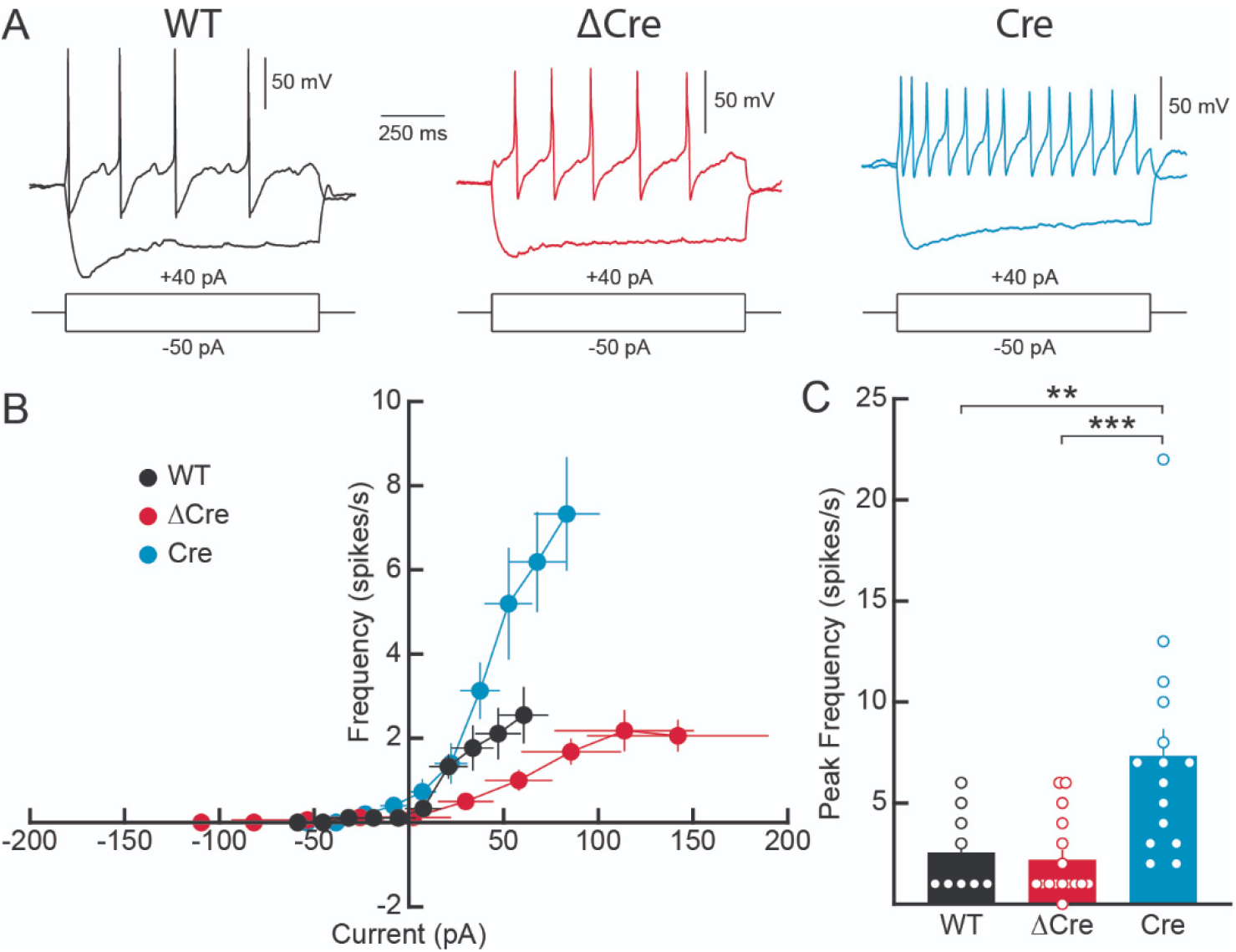
Genetic knockout of SUR1 subunits increases DRG excitability. A) Example traces of responses to hyperpolarizing and depolarizing current application in WT (black traces), ΔCre control (red traces) and Cre knockdown (blue traces). B) Mean F/I curves of the three groups. The equation for the linear portion of each FI curve was determined to be: WT (n = 9), y = 0.039X + 0.28; ΔCre (n = 16), y = 0.019X + 0.23; Cre (n = 14), y = 0.098X – 0.52. The slopes were determined to be significantly different based on a simple linear regression (F = 11.68, p < 0.0001). C) Peak frequency analysis during the current step applications. The Cre peak frequency was measured to be singificantly higher than both WT (p = 0.0083) and ΔCre (p = 0.0008) as determined by a one-way ANOVA and Tukey’s multiple comparison post-hoc analysis. Error bars are standard error of the means.

**Figure 3.**
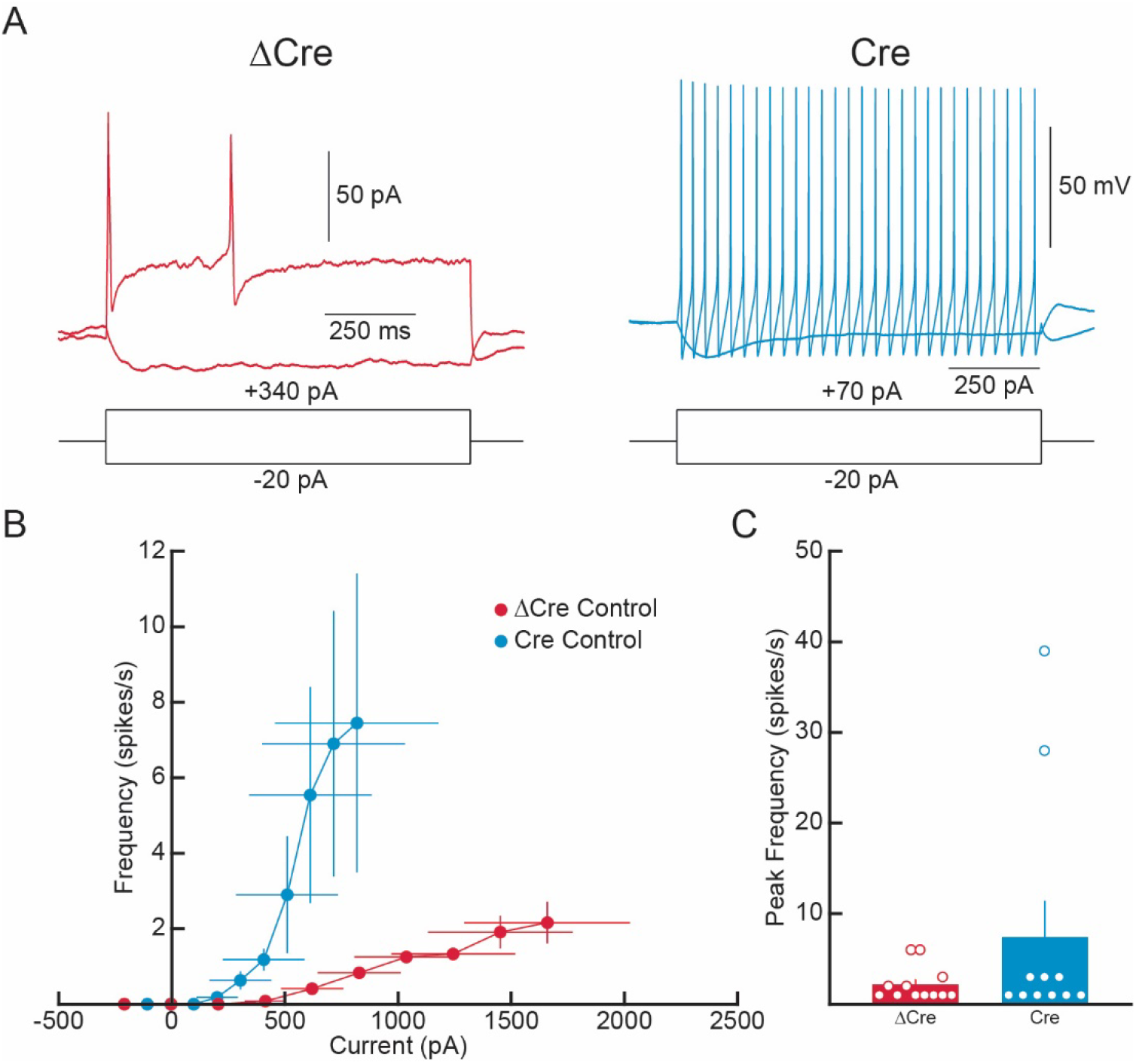
Knockout of K_IR_6.2 increases DRG neuron excitability. A) Example traces of current step applications in KIR6.2-floxed DRG neurons injected with ΔCre and Cre. The Cre example is used to exemplify one of the few cells that were found to be dramatically changed. B) Mean F/I curves of the two conditions. The slopes of the F/I curve were found to be significantly different (F = 9.066, p = 0.0032). Linear fits were as follows: ΔCre Control: y = 0.001564x – 0.4551 and Cre Control: y = 0.01643x – 4.942. C) Peak frequencies achieved during the current step applications were not found to be singificantly different. While DRG from some of the Cre injected animals appeared to have a higher peak frequency, collectively they were not singificantly different from controls(p = 0.2145).

### Potassium flux is inhibited by conditional deletion of SUR1 or Kir6.2 after pharmacological stimulation

We conducted potassium flux assays, using cultured mouse DRG, with and without stimulation from the K_ATP_ channel SUR1-subtype agonist, Diazoxide. Kir6 and SUR subunits both contain endoplasmic reticulum retention motifs that prevent the surface expression of either subunit alone (Zerangue et al., 1999), and therefore loss of one subunit should lead to lack of channel expression/function of the other subunit. Diazoxide was used to stimulate K_ATP_ channels in DRG due to several studies providing evidence that diazoxide decreases DRG excitability in acute and pain models (Kawano, Zoga, Gemes, et al., 2009; Sarantopoulos et al., 2003; Xu et al., 2024; Zoga et al., 2010). A red-fluorescent protein (RFP) marker allowed for separation of DRG based on uptake of control virus (ΔCre) or Cre-virus in Kir6.2 or SUR1 cKO mice (Figure 4A-C). In neurons that did not have RFP signal above threshold levels (-RFP), we see that diazoxide significantly increased fluorescent signal, corresponding to increased cation flux, in control vector mice. This comparison is also true in Kir6.2 and SUR1 cKO mice, neurons that received diazoxide have significantly higher fluorescence values than thallium alone (Figure 8D). Interestingly, Kir6.2-cKO mice have a lower fluorescence after application of thallium alone compared to the control virus-treated mice (Figure 4D). In neurons with a RFP signal above threshold, diazoxide also significantly increased fluorescent signal in control vector mice but did not increase fluorescence in Kir6.2 or SUR1 cKO mice (Figure 4E). In the cells from Kir6.2 cKO mice, diazoxide application significantly decreased potassium flux compared to cells without diazoxide application (Figure 4E).

**Figure 4.**
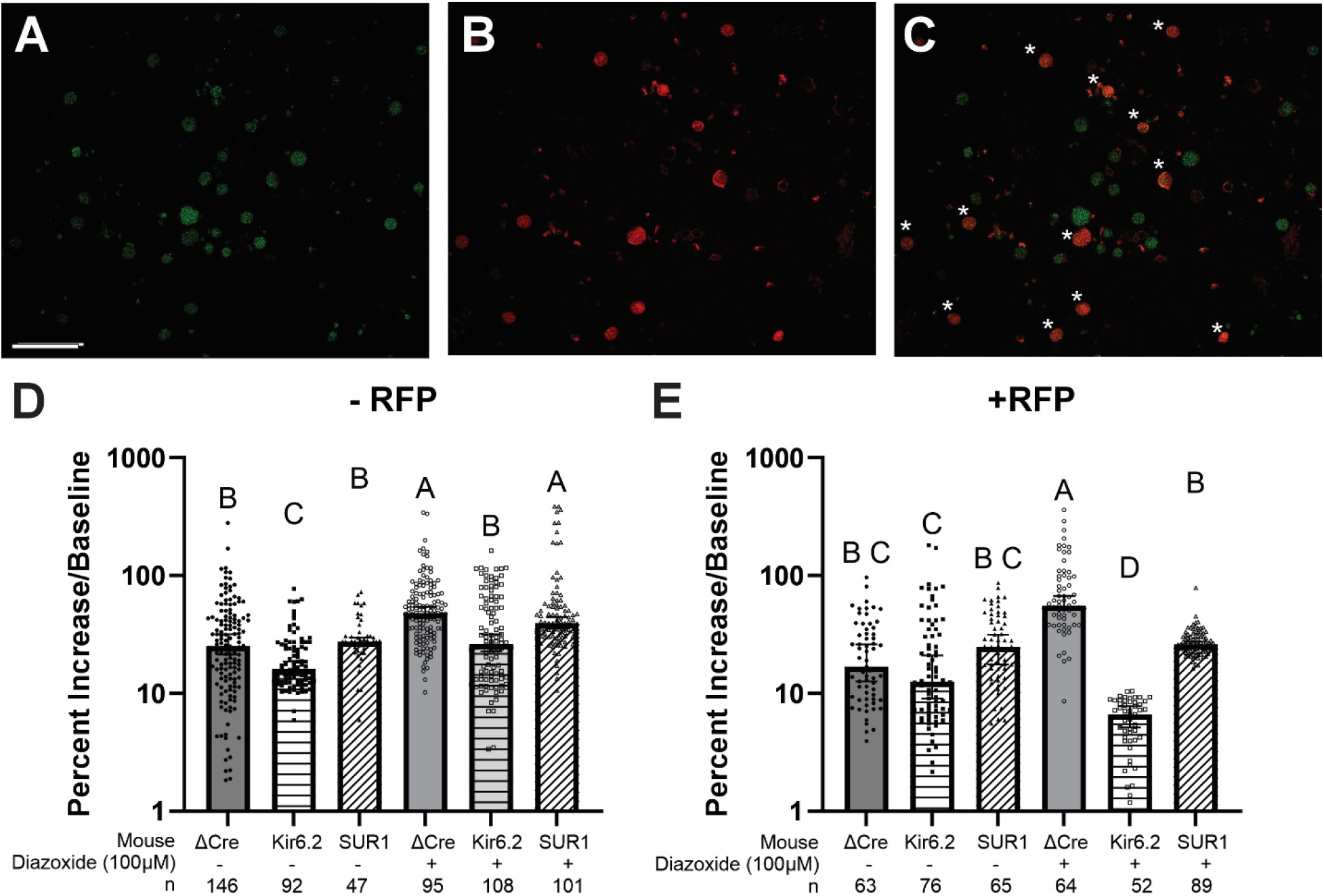
Loss of SUR1 and Kir6.2 in mouse dorsal root ganglia (DRG) reduces sensitivity to diazoxide. Kir6.2 or SUR1 floxed mice were inoculated with AAV9-HSyn-ΔCre-RFP (ΔCre) or AAV9-HSyn-Cre-RFP (Kir6.2 or SUR1) via intrathecal injection. DRG were cultured overnight and loaded with a thallium indicating dye to measure potassium flux. Photomicrographs of (A) mouse DRG loaded with thallium dye, (B) cells positive for red fluorescent protein (RFP) and (C) overlay between A and B. Cells marked with asterisks (*) positive for dye loading and RFP signal. (D) Percent increase in fluorescence of individual cells after addition of thallium with or without diazoxide (100uM) without RFP signal. (E) Percent increase in fluorescence after addition of thallium with or without diazoxide in cells with RFP signal. Data taken from 5-8 mice per group over five minutes post thallium addition. Images taken from Leica SP8 STED confocal at 20x magnification. Scalebar in (A) is 100uM. Statistical analysis done by one-way ANOVA with Tukey-post hoc. Groups labeled with condensed letter format where groups denoted with the same letter are not statistically significantly different.

### Decreased morphine antinociception after conditional downregulation of SUR1 or Kir6.2

In previous studies, loss of K_ATP_ channel expression or function decreased opioid signaling and antinociception (Cunha et al., 2010; Fisher, Johnson, et al., 2019; Sakamaki et al., 2021). The mechanical (Figure 5A) and heat thermal thresholds (Figure 5B) of Kir6.2 and SUR1 flox mice after conditional deletion (AAV9-HSyn-Cre-RFP) were similar to Kir6.2/SUR1 cKO mice with (AAV9-HSyn-ΔCre-RFP, ΔCre) and without (No injection) intrathecal injections. Escalating morphine dose response curves indicate that Kir6.2 and SUR1 flox mice injected with AAV9-HSyn-Cre-RFP, have lower mechanical thresholds, compared to non-injected and ΔCre (Figure 5C). Thermal thresholds across morphine doses were comparable across all treatment groups (Figure 5D). Area under the curve calculations for the morphine dose response data indicate a significant difference in morphine antinociception with mechanical threshold testing in Kir6.2 and SUR1 flox mice (Figure 5E), which was not seen with thermal thresholds (Figure 5F). Validation of *Kcnj11* (protein: Kir6.2) and *Abcc8* (protein: SUR1) downregulation was seen in spinal cords collected from animals in behavioral assays and imaging studies (Supplemental Figure 1).

**Figure 5.**
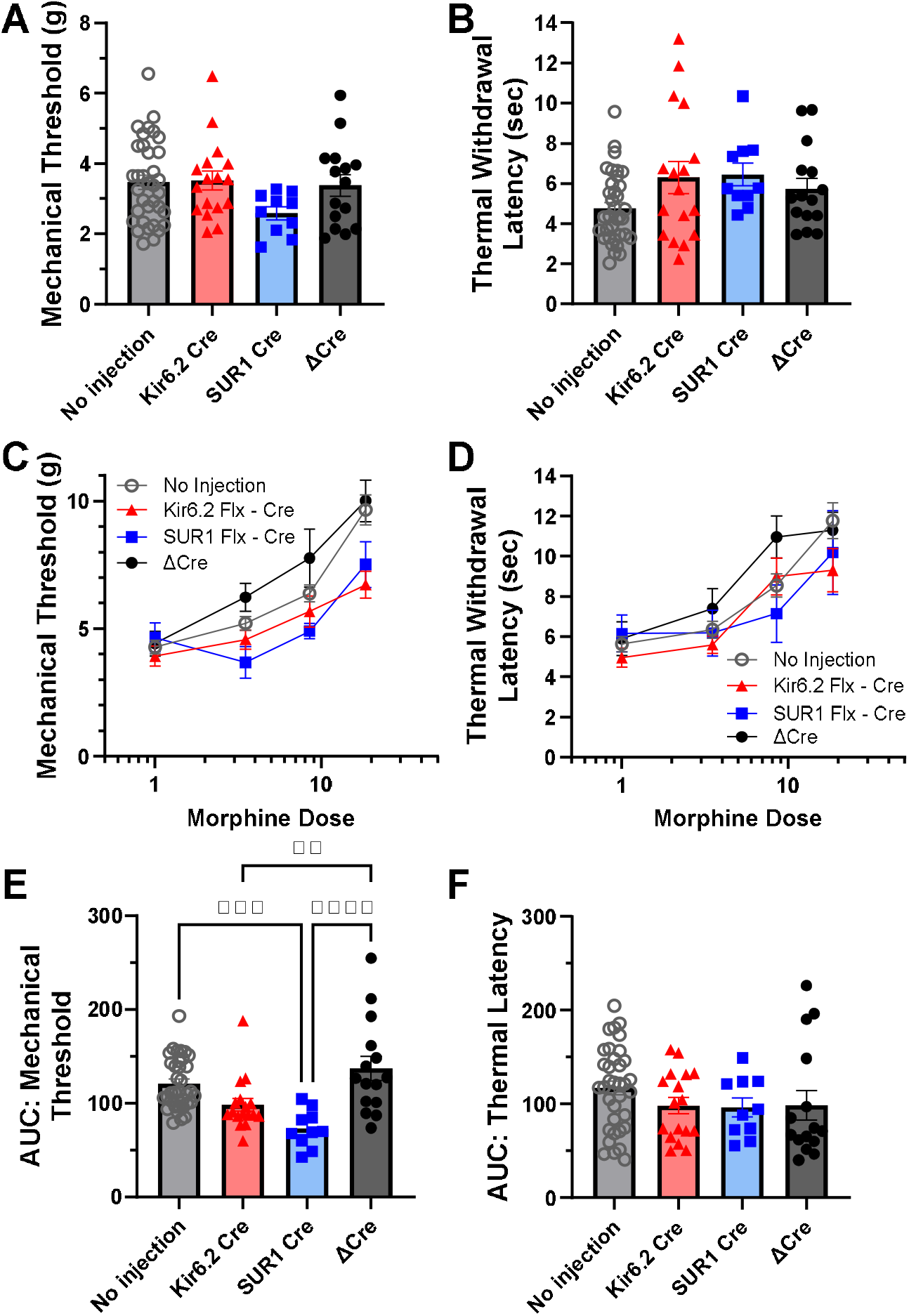
Loss of Kir6.2 or SUR1 reduces morphine antinociception in mice. Kir6.2 or SUR1 floxed mice were inoculated with AAV9-HSyn-ΔCre-RFP (ΔCre) or AAV9-HSyn-Cre-RFP (Kir6.2 or SUR1) via intrathecal injection. Kir6.2 or SUR1 floxed without injection were also compared. Baseline mechanical thresholds (A) or thermal thresholds (B) were not different across treatment group. (C) Cumulative morphine dose response curves for mechanical thresholds and (D) thermal thresholds. (E) Area under the dose response curve in (C) indicates that loss of SUR1 and Kir6.2 decrease morphine antinociception. (F) Area under the dose response curve in (D), thermal latencies are not affected by loss of Kir6.2 or SUR1.

### Chronic opioid exposure decreases RNA levels of Abcc8 and Kcnj11 in mouse DRG

Previous studies have implicated a loss in K_ATP_ channel activity and expression in after chronic opioid administration (Salman et al., 2014) and administration of drugs that open K_ATP_ channels have the ability to decrease withdrawal and attenuate opioid tolerance (Doucette et al., 2023). We wished to determine if genetic changes were restricted to the spinal cord or DRG, and if chronic opioid exposure affected each subunit in an equal manner. *Abcc8* and *Kcnj11* RNA were detected in both mouse DRG (Figure 6A,B) and spinal cord (Figure 6C). A significant reduction in cells positive for *Kcnj11* RNA signal was observed from mice treated with morphine for five days (Figure 6B). We also observed a decrease in the number of DRG cells positive for both *Abcc8* and *Kcnj11 in morphine treated mice*. We did not observe a significant change in the percentage of cells positive for either *Abcc8* or *Kncj11* in the spinal cord (Figure 6C). Cells were also scored according to number of punta visible, which indicated that *Kcnj11* signal was significantly altered in DRG morphine treated mice (Table 2), notably where the lowest score was significantly elevated compared to saline treated animals. No changes were detected in the spinal cord with regards to scoring of punta within cells across treatment groups. There was also a significant decrease in the percent of cells that labeled for either *Abcc8* and *Kcnj11* in morphine treated mice. In DRG, the percentage of cells that did not label for either probe increased from 88.8 ± 1.76% in saline treated mice to 94.7 ± 0.62 % in morphine treated mice (Unpaired t-test: P = 0.0017), and in spinal cord this value also increased from 65.38±5.45% in saline treated mice to 79.10 ± 1.81% in morphine treated mice (Unpaired t-test: P = 0.015).

**Figure 6.**
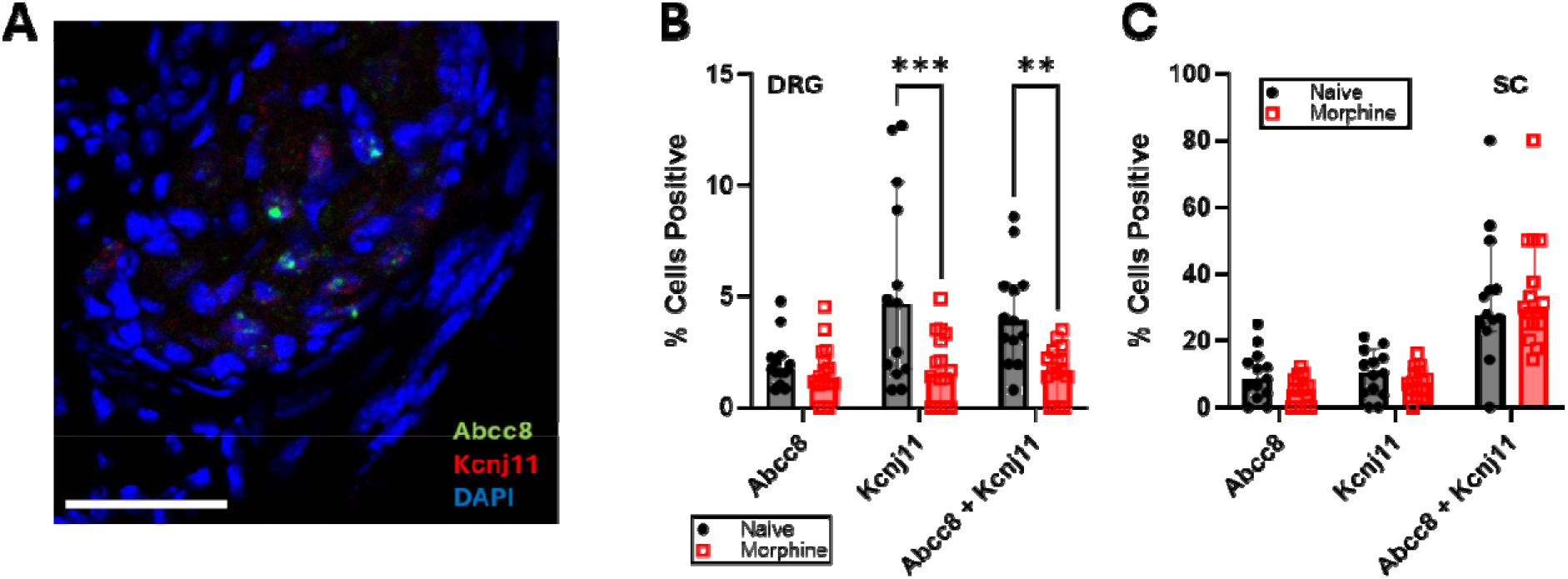
RNA expression of K_ATP_ channel subunits, Kir6.2 and SUR1 are decreased in mouse dorsal root ganglia (DRG) after chronic morphine exposure. (A) Representative image of mouse DRG labeled with RNAscope *in situ* hybridization for *Abcc8* (green) and *Kcnj11* (red) mRNA and co-stained with DAPI (blue). Scalebar = 100μM. (B) Mice treated with morphine (15mg/kg, twice daily x 5 days) have significantly less expression of *Kcnj11* and co-expression of *Abcc8* and *Kcnj11* in DRG (Mann-Whitney test to compare ranks, P = 0.0193 and P = 0.0008, respectively). (C) *Abcc8* and *Kcnj11* expression were not significantly decreased in spinal cord of mice treated with morphine compared to naïve animals.

**Table 2.** Percent of mouse DRG with RNAScope® Scores positive for probes Abcc8 and/or Kcnj11 mRNA. Scoring criteria were, 1: 1-3 dots/cell, 2: 4-9 dots/cell, 3: 10-15 dots/cell and/or <10% dots in clusters, 4: >15 dots/cell and/or >10% dots in clusters. *P = 0.0223 Mann-Whitney test to compare ranks between Naïve (n = 13) and morphine (n = 16) treated animals. (Kcnj11: Holm-Sidak method to correct for multiple comparisons).

|  | Naïve: Abcc8 | Morphine: Abcc8 | Naïve: Kcnj11 | Morphine: Kcnj11 |
| --- | --- | --- | --- | --- |
| Score 1 | 26.99 ± 7.68 | 29.90 ± 10.77 | 20.83 ± 8.67 | 60.24 ± 11.34 * |
| Score 2 | 43.74 ± 7.80 | 38.33 ± 9.53 | 36.46 ± 8.68 | 19.62 ± 6.11 |
| Score 3 | 23.12 ± 5.60 | 27.08 ± 8.69 | 23.44 ± 7.48 | 11.35 ± 4.87 |
| Score 4 | 6.15 ± 3.31 | 4.69 ± 3.40 | 13.02 ± 5.64 | 1.10 ± 1.10 |

**Table 3.** Percent of mouse spinal cord cells with RNAScope® Scores positive for probes Abcc8 and/or Kcnj11 mRNA. Scoring criteria were, 0: No staining, 1: 1-3 dots/cell, 2: 4-9 dots/cell, 3: 10-15 dots/cell and/or <10% dots in clusters, 4: >15 dots/cell and/or >10% dots in clusters.

|  | Naïve: Abcc8 | Morphine: Abcc8 | Naïve: Kcnj11 | Morphine: Kcnj11 |
| --- | --- | --- | --- | --- |
| Score 1 | 29.76 ± 8.37 | 24.09 ± 6.54 | 80.05 ± 10.03 | 76.48 ± 7.68 |
| Score 2 | 39.48 ± 5.99 | 33.20 ± 5.17 | 9.34 ± 5.14 | 13.88 ± 4.50 |
| Score 3 | 25.03 ± 6.30 | 34.04 ± 7.19 | 2.91 ± 2.56 | 3.39 ± 1.91 |
| Score 4 | 5.73 ± 2.69 | 8.66 ± 3.03 | 0.00 ± 0.00 | 0.00 ± 0.00 |

## Discussion

In this study we determined that the loss of either Kir6.2 or SUR1 affected peripheral neuron excitability, morphine antinociception, and expression after chronic morphine administration to a similar degree. Electrophysiological recordings from DRG neurons determined that K_ATP_ channel agonists and antagonists decreased and increased action potential firing, respectively. Neuronal recordings in mice after administration of viral Cre-recombinase in Kir6.2^flox/flox^ and SUR1 ^flox/flox^ mice significantly increased firing activity compared to knockdown controls. Baseline sensitivity to mechanical or heat stimulation was not changed in Kir6.2 or SUR1 conditional knockdown, but morphine-induced antinociception was attenuated. In situ hybridization data indicated that cells in the DRG or spinal cord co-expressed mRNA transcripts for *Abbc8* and *Kcnj11*, but *Abcc8* expression was more affected after chronic exposure to morphine. Overall, it appears that the loss of SUR1 did not affect output measures in this study much differently than loss of Kir6.2, indicating that despite expression pattern differences seen with sequencing data and immunohistochemistry(Zoga et al., 2010), SUR1 and Kir6.2 most likely form heteromeric complexes to influence neuronal excitability under naïve conditions.

The electrophysiological recordings from DRG neurons using pharmacological and genetic approaches demonstrated that peripheral sensory neurons could be modulated by increasing or decreasing the activity of Kir6.2 or SUR1 subunits. Glibenclamide, known to inhibit SUR1 and promote exocytosis of insulin granules in pancreatic βcells, induced depolarization and increased action potential firing. Glibenclamide has also been shown to depolarize cardiac and pelvic ganglionic neurons in other *in vitro* systems (Hogg & Adams, 2001; Kyu-Sang et al., 2002). SUR1 appears to be the predominant K_ATP_ channel regulatory subunit expressed in DRG, as preincubation with a SUR1-antibody prevents the blocking effect of glibenclamide on single K_ATP_ channel opening in excised membrane patches (Zoga et al., 2010). The K_ATP_ channel agonists, diazoxide and CL-705G decreased action potentials upon increasing current stimulation and decreased the resting membrane potential. This agrees with studies utilizing other neuron types, as membrane hyperpolarization and loss of spontaneous action potentials are seen after diazoxide incubation in whole-cell patch clamp recordings in cultured hippocampal neurons (Liu et al., 2010). Diazoxide exhibits some cross activation of SUR2B-containing channels which are typically found in smooth muscle (Coetzee, 2013), but DRG currents activated by diazoxide can be completely blocked by low concentration of glibenclamide (Du et al., 2011).

The increase in DRG neuron action potential firing and loss of potassium flux was enhanced by genetic reduction of SUR1 or Kir6.2 subunits, suggesting that attenuating the activity of either subunit greatly changes neuronal excitability. The Kir6.2 subunit has been shown to be able to form a functional channel in the absence of SUR1 (John et al., 1998), but only under experimental conditions where the C-terminal tail has been removed to by bypass trafficking checkpoints, letting the subunit reach the plasma membrane without the regulatory sulfonylurea receptor. Similarly, the purified SUR1 subunits have been shown to form tetramers(de Wet et al., 2007), under optimal experimental conditions. In this study, the loss of Kir6.2 resulted in decreased sensitivity to diazoxide in flux studies, even though diazoxide interacts with nucleotide binding folds on the SUR subunits (Shyng et al., 1997). mRNA expression of *Kcnj11* was significantly reduced in Kir6.2 recombinant animals but was also decreased in SUR1-recombinant animals. In a previous study, intrathecal delivery of a viral vector used to promote upregulation of Kir6.2 (Ad-m-Kcnj11) increased expression of both *Kcnj11* and *Abcc8* in the spinal cord (Fisher et al., 2024), suggesting that expression of the pore-forming subunit may be interdependent with the regulatory subunit. Although the theoretical possibility exists for Kir6.2 and SUR1 to function individually without each other, the electrophysiological and imaging data indicate that in vivo these channels may not typically operate in this fashion.

In behavioral studies, the global loss of SUR1 subunits has been shown to be associated with increased baseline mechanical sensitivity (Luu et al., 2019) and reduce opioid-induced antinociception (Sakamaki et al., 2021). In the current study, we did not see a significant reduction in baseline mechanical or thermal sensitivity, which is consistent with a previous study, that showed that these values were largely consistent after recombination in neurons SUR1 flox mice (Fisher, Johnson, et al., 2019). In Kir6.2 global knock out mice, sensory nerve conduction velocities were found decreased compared to age-matched controls at 24 weeks, and thermal paw withdrawal latencies to heat was significantly elevated (Nakai-Shimoda et al., 2022). Baseline mechanical thresholds and heat withdrawal latencies were not greatly impacted in Kir6.2 flox mice used in this study. Previous work has shown that Kir6.2 knockout mice initially show only a very mild impairment in glucose tolerance, and hyperglycemia and glucose intolerance occur only in obese mice after 60 weeks (Seino et al., 2000), so the deficiencies may not have appeared in our study due to the younger age of mice or because the knockdown was restricted to neurons in our study. Loss of Kir6.2 subunits also decreased morphine antinociception with mechanical, but not thermal stimulation, which has been seen previously with SUR1 (Fisher, Johnson, et al., 2019).

This study did not investigate tissues or behavioral outputs from injury or disease models. The dysregulation of K_ATP_ channels has been implicated in various neurodegenerative diseases. In contexts such as Alzheimer’s disease, hyperexcitability of neurons is a key feature, where altered K_ATP_ channel activity influences cellular excitability and may promote neurodegeneration (Liu et al., 2010). Prior studies have determined that loss of K_ATP_ channel subunits occurs in nerve injury models (Luu et al., 2019; Zoga et al., 2010) suggesting that these channels contribute to neuroprotective functions in DRG, which are decreased after physical insult. This study also focused specifically on neuronal expression and function of K_ATP_ channel subunits.

Given the diverging roles of K_ATP_ channel expression in glia, further studies would be necessary to determine the impact of altered K_ATP_ channel subunit expression in satellite cells, microglia, and astrocytes. Comprehensive interaction data modeling suggests that Kir6.2 or SUR1 can form protein-protein interactions with alternative proteins (Szklarczyk et al., 2023), including TRPM4, a non-selective monovalent cation channel consisting of four SUR1 subunits that bind to four TRPM4 subunits (Mehta et al., 2015), but this has been debated(Sala-Rabanal et al., 2012). The SUR1-TRPM4 channel is minimally expressed in the healthy brain but reportedly becomes pathologically upregulated de novo in neurons, glia and endothelial cells after injury in rodent models (Tsymbalyuk et al., 2021). These changes may be restricted to the spinal cord, as a single cell RNAseq study analyzing DRG after sciatic nerve injury suggests mostly neuronal of pericyte expression of *Abcc8* (Jager et al., 2022). Future studies could investigate potential interactions in non-neuronal cells or if these interactions change in chronic pain models.

In our studies, we determined that chronic morphine exposure decreased expression of *Kcnj11* and *Abcc8* mRNA in spinal cord and DRG using in situ hybridization. These data disagree with in vitro studies (Salman et al., 2014) but are in alignment with previous in vivo studies, indicating *Kcnj11* and *Abcc8* transcripts are downregulated in DRG after chronic opioid exposure (Fisher, Janecek, et al., 2019). Moreover, the in-situ hybridization data of *Abcc8* and *Kcnj11* in spinal cord and DRG indicate that expression of each subunit varies but overall agree with sequencing data suggesting low expression in DRG (3-10%; (Bhuiyan et al., 2024; Krauter et al., 2025).

Spinal cord sequencing data suggests much lower expression of *ABCC8* spread across various cell types in humans (Yadav et al., 2023). In comparison with the in-situ hybridization data presented in the current study, the co-localization of *Abcc8* and *Kcnj11* in the spinal cord was minimal. The mismatch between reported RNAseq data and in situ hybridization could be due to RNA degradation, splice variants (Schmid et al., 2012), and/or differences in assay sensitivity.

Targeting K_ATP_ channel subunits modulate neuronal excitability, regardless of subunit. Their involvement in both normal physiology and pathological states emphasizes their potential utility as therapeutic targets in neurodegenerative diseases and pain management. Although several K_ATP_ channel targeting drugs exist, none have been utilized in the clinic so far for analgesia or opioid use disorders. Pharmacological strategies to target sulfonylurea subunits have been around for decades, with largely high tolerability, despite lack of specificity in most cases(Jahangir & Terzic, 2005). Several studies suggest that K_ATP_ channel agonists are beneficial for several nervous system disorders, including pain. For example, cromakalim has been shown to reduce neuropathic pain (Wu et al., 2011) although it’s activity at SUR2 and SUR1 activate both smooth muscle and cardiac muscle K_ATP_ channels. It’s possible that by chemically targeting only Kir6.2-containing channels (Gando et al., 2023), such as those in the peripheral nervous system, we could be able to target tissues involved in disrupted metabolism and somatosensation. Continued research into K_ATP_-subunit targeting strategies may uncover novel insights into the intricate workings of neuronal excitability and its implications for brain and nervous system health.

## Acknowledgements

This work was supported by the National Institutes of Health, R01DA051876 to A.H.K., W.A.C, and S.C.S.

