## Supplemental Figures for "Loss of SUR1 and Kir6.2 ATP-sensitive potassium channel subunits produces comparable neuronal excitability and morphine antinociception"

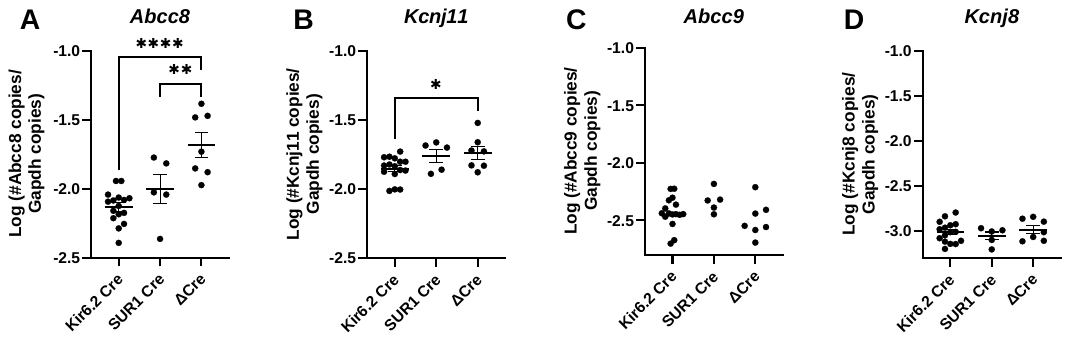

**Supplemental Figure 1.** Quantitative PCR for K_ATP_ channel subunits in spinal cord after intrathecal injection of AAV9-HSyn-ΔCre-RFP (ΔCre) or AAV9-HSyn-Cre-RFP in Kir6.2 or SUR1 floxed mice. (A) The number of copies for Abcc8 (protein: SUR1), was significantly decreased in Kir6.2 (P < 0.001) and SUR1 (P = 0.0078) flox animals after Cre-recombination compared to Kir6.2 and SUR1 flox animals with the control virus (ΔCre). (B) The number of Kcnj11 copies was significantly decreased in Kir6.2 flox mice after recombination compared to control mice (P = 0.034). Copies of other K_ATP_ channel subunits including (C) Abcc9 (protein: SUR2) or (D) Kcnj8 (protein: Kir6.1, typically not found in neurons) were not significantly different across treatment groups. Analysis by one-way ANOVA with Dunnett’s post-hoc test.
